# Morphological and genetic heterogeneity in *Aedes aegypti* (Diptera: Culicidae) populations across diverse landscapes in West Africa

**DOI:** 10.1101/2025.08.06.668854

**Authors:** Eric Agboli, Julien Z B Zahouli, Felix Gregor Sauer, Aboubacar Sombie, Yasmine N Biré, Claver N Adjobi, Daniel Cadar, Balázs Horváth, Alexandru Tomazatos, Jonas Schmidt-Chanasit, Renke Lühken, Gábor Endre Tóth, Athanase Badolo, Hanna Jöst

**Affiliations:** Bernhard Nocht Institute for Tropical Medicine, WHO Collaborating Centre for Arbovirus and Haemorrhagic Fever Reference and Research, Hamburg, Germany; University of Health and Allied Sciences, School of Basic and Biomedical Sciences, Ho, Ghana; Centre d’Entomologie Médicale et Vétérinaire, Université Alassane Ouattara, Bouake, Côte d’Ivoire; Centre Suisse de Recherches Scientifiques en Côte d’Ivoire, Abidjan, Côte d’Ivoire; Faculty of Mathematics, Informatics and Natural Sciences, Universität Hamburg, Hamburg, Germany; Laboratory of Fundamental and Applied Entomology, Université Joseph Ki-Zerbo, Ouagadougou, Burkina Faso

**Keywords:** *Aedes aegypti*, Morphology, Abdominal scaling, Morphometry, Wing shape, Genetics, arbovirus vector, Africa, Burkina Faso, Côte d’Ivoire, Ghana

## Abstract

Native to sub-Saharan Africa, *Aedes aegypti* has spread across the globe and is now one of the most significant vectors of arboviruses worldwide. However, data on the ranges of its populations remain sparse, and the genetic variability and ecological adaptability in West Africa are still poorly understood. In this study, we characterized the morphological and genetic diversity of *Ae. aegypti* across four landscape types (urban, peri-urban, rural, and sylvatic sites) in three West African countries (Burkina Faso, Côte d’Ivoire, and Ghana). *Ae. aegypti* exhibited significant variation in abdominal scaling patterns across countries and landscape types, with the sylvatic and urban populations in Burkina Faso displaying the highest proportions of white scales (>50% white scales), while black scales predominated among those from Côte d’Ivoire and Ghana (>80% black scales). Wing shape displayed limited differentiation between the countries, landscape types, and genetic clusters. Bayesian analysis indicated high gene flow among populations, with notable outliers observed in sylvatic sites from Burkina Faso and admixture patterns suggesting possible human-mediated dispersal. Additionally, two major mitochondrial lineages, clades A and B, were identified. Most samples were categorized under clade B, showing no evidence of clustering by country or landscape type. In contrast, clade A comprised primarily sylvatic specimens from Burkina Faso and a single urban individual from Côte d’Ivoire. These findings highlight the complex interplay of genetic, environmental, and ecological factors shaping the variations in *Ae. aegypti* populations in West Africa. They provide insights into the phenotypic and genetic diversity of *Ae. aegypti*, offering valuable implications for understanding arbovirus transmission dynamics and formulating targeted interventions against arboviral diseases.

## Introduction

*Aedes (Stegomyia) aegypti* is one of the most important vectors for arboviruses worldwide. In West Africa, it is known to transmit dengue virus, yellow fever virus, chikungunya virus, and Zika virus (Agboli et al. 2021). These pathogens pose a significant public and global health threat, with over 8 millions of people affected each year (Weetman et al. 2018).

The evolutionary origin of *Ae. aegypti* lies in sub-Saharan Africa from where it has spread global and has invaded all five continents (Africa, America, Asia, Europe, and Oceania). The ancestral form was likely a generalist tree-hole breeder, feeding on animal hosts (Brown et al. 2014). This form was given the subspecies rank *Ae. aegypti formosus* (Aaf), which is much darker than the urban or domestic form feeding on humans (*Ae. aegypti aegypti*, Aaa), and was considered to never has pale or white scales on the first abdominal tergite (Mattingly 1957). Due to inconsistency of morphological features with ecological and behavioral traits, the accuracy and utility of subspecies designation have come under scrutiny (McBride et al. 2014; Powell and Tabachnick 2013). Recently, Harbach and Wilkerson (2023) raised a concern regarding the unsupported validity of mosquito subspecies and their exclusion from culicid classification.

Population genetic studies on *Ae. aegypti* in Africa are limited, posing a challenge for understanding genetic variability, adaptation, gene flow over time, and their associated epidemiological implications. A worldwide investigation involving genetic diversity across 12 microsatellite loci of *Ae. aegypti* showed that the two subspecies Aaf and Aaa are genetically distinct entities (Gloria-Soria et al. 2016). Genetic evidence suggests that *Ae. aegypti* moved to continental Africa less than 85,000 years ago, with its origin traced to the southwestern Indian Ocean (Soghigian et al. 2020). Using genome-wide single nucleotide polymorphisms (SNPs), West Africa was identified as the source of the New World’s invasion (400-500 years ago) of *Ae. aegypti* and later Asia (Kotsakiozi et al. 2018). This finding aligns closely with the documented history of the 16^th^ century transatlantic slave trade between Africa and the Americas (Kotsakiozi et al. 2018).

Mitochondrial DNA (mtDNA) genes have been widely utilized to investigate the genetic relationships of *Aedes* mosquitoes, though most studies rely on analyzing only a short segment of the mtDNA (Abuelmaali et al. 2022; Hounkanrin et al. 2023; Moore et al. 2013; Paupy et al. 2008; Thia et al. 2024). These genes serve as markers for studying ancestry and population dynamics due to their uniparental (maternal) inheritance, lack of recombination and high mutation rates (Avise 1995). However, population genetic studies targeting certain regions of the mtDNA such as cytochrome c oxidase I and NADH dehydrogenase subunit 5 genes have revealed some limitations, as the observed genetic variation may not be sufficient to reliably assign mosquitoes to distinct haplogroups (Zé-Zé et al. 2020). The advent of next-generation sequencing (NGS) has enabled the analysis of thousands to millions of SNPs, providing deeper insights into populations’ structure and ancestry.

Geometric morphometrics uses the coordinates of homologous landmarks or semi-landmarks on biological structures to examine their shape and size variation (Klingenberg 2010). In mosquitoes, the wings are usually used for geometric morphometric analysis, as the anatomical junctions of the wing veins provide an easy target for landmark collection (Lorenz et al. 2017). Wing geometric morphometrics (WGM) has demonstrated its effectiveness in accurately distinguishing mosquito species, making it a valuable complement to traditional methods of morphological and molecular identification (Brown et al. 2011; Gloria-Soria et al. 2016; Jeon et al. 2024; Lewis 1945; Paupy et al. 2010; Sauer et al. 2020). Wing shape analyses serve as a valuable tool for studying mosquito population dynamics (Wilk-da-Silva et al. 2018). Moreover, wing morphology can be influenced by biotic and abiotic conditions of breeding habitats through carry-over effects from the immature stages to adulthood (Evans et al. 2018; Jirakanjanakit et al. 2007; Ouédraogo et al. 2022; Phanitchat et al. 2019; Roux et al. 2015). In Africa, WGM studies are scarce, especially with native *Ae. aegypti* mosquitoes. To the best of our knowledge, WGM of *Aedes* mosquitoes was investigated only in Benin (Hounkanrin et al. 2023).

The combination of population genetics and WGM has proven to be a powerful tool in studying population structuring in mosquitoes (Carvajal, Amalin, and Watanabe 2021; Dujardin 2008; Laojun, Changbunjong, and Chaiphongpachara 2024). In *An. cruzii*, wing shape and cytochrome c oxidase I gene analyses demonstrated altitudinal population structuring (Lorenz et al. 2014). Wing shape analysis successfully distinguished between *Anopheles* populations, but was less effective for *Ae. aegypti* populations compared to microsatellite analysis (Vidal and Suesdek 2012). However, wing morphometrics revealed population structuring in *Ae. aegypti* related to urbanization levels (Wilk-da-Silva et al. 2018). These studies suggest that combining wing morphometrics and genetic analysis can be valuable for assessing mosquito population structure, although the effectiveness can vary depending on the species and environmental factors. Additionally, by integrating genetic information with morphometric and morphological data, a more comprehensive understanding of factors influencing mosquito populations, such as migration patterns, ecological preferences, and the impact of control interventions can be achieved.

This study investigated *Ae. aegypti* morphological diversity and population genetic structure in three West African countries, namely, Burkina Faso, Côte d’Ivoire and Ghana. By analyzing the genetic variation and wing shape differences among these *Ae. aegypti* populations, we aimed to enhance our understanding of their population structure and migration patterns.

## Materials and Methods

### Study sites

The study was conducted in three neighboring countries, Burkina Faso, Côte d’Ivoire and Ghana, located in West Africa (Figure 1).

**Figure 1:**
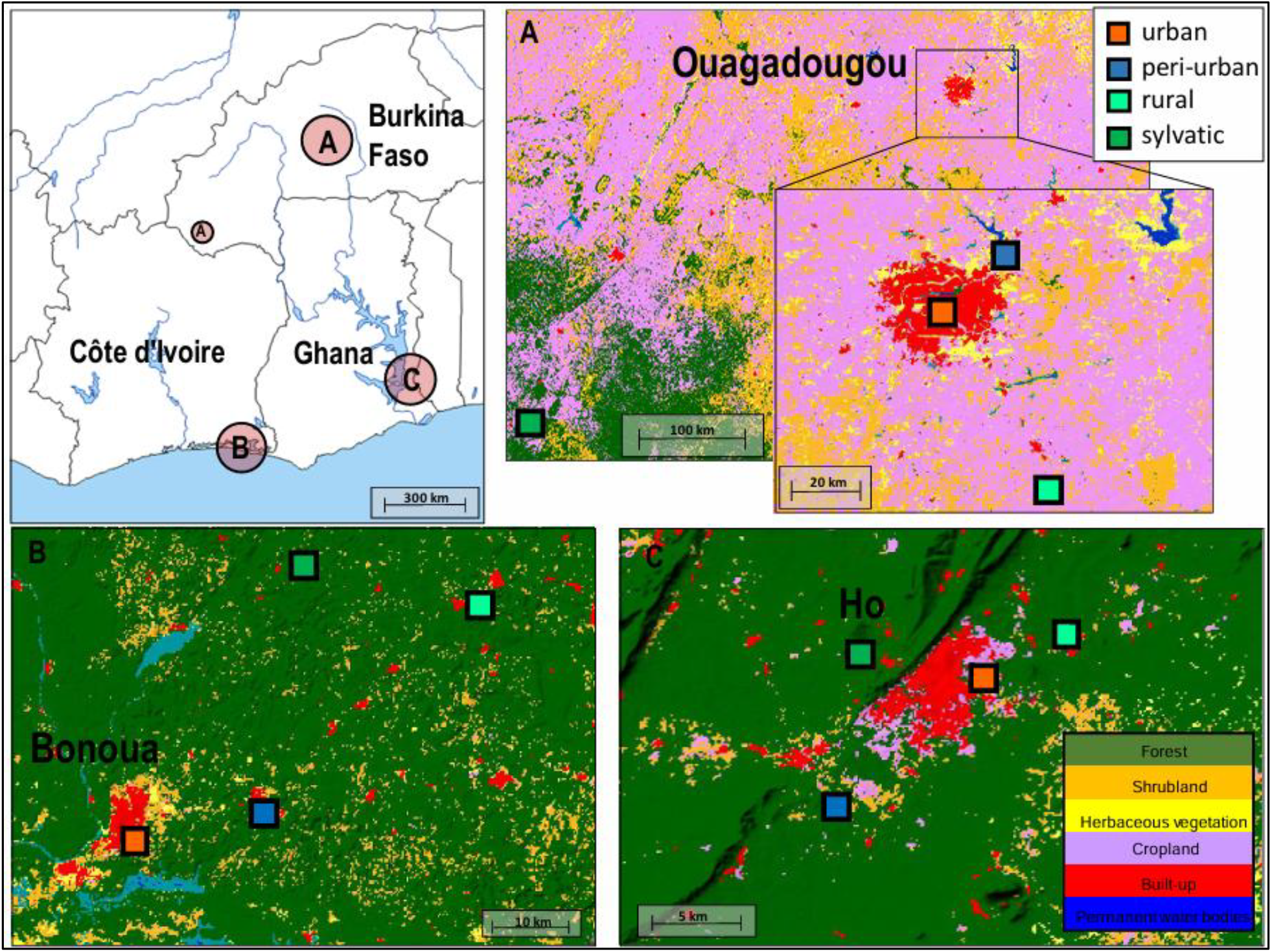
Study sites showing ecological features. The study areas for Burkina Faso, Côte d’Ivoire, and Ghana are (A) Ouagadougou, (B) Bonoua, and (C) Ho, respectively. Colours represent the land cover types; green (forest), orange (shrubland), yellow (herbaceous vegetation), periwinkle (cropland), red (built-up), and blue (permanent water bodies). © = Rural site in Burkina Faso not part of the city of Ouagadougou (Source of maps: Land Cover Viewer, Nazka Maps (https://www.nazka.be/en/realisations/landcoverviewer))

In Burkina Faso, *Aedes* mosquitoes were collected in urban Ouagadougou and its surrounding areas. The region has a hot semi-arid climate with distinct dry and wet seasons and consistently high temperatures throughout the year. The dry season lasts from November to March, while the rainy season occurs from May to October, with peak rainfall in August and the highest temperatures, exceeding 40°C, in March and April.

In Côte d’Ivoire, *Aedes* samples were collected in the region of Bonoua, about 50 km east of Abidjan. The local climate is warm and humid. It is distinguishable by rainfall and has two rainy and two dry seasons: the long rainy season lasts from April to July whilst the short rainy season runs from September to November, both being separated by the long and short dry seasons.

In Ghana, *Aedes* samples were collected in the Ho region, located in the southeastern part of the country near the border with Togo. The region experiences a tropical climate characterized by two main seasons: the wet and dry seasons. The wet season typically runs from April to October whilst the dry season lasts from November to March. The climate is influenced by its proximity to the equator and its low elevations.

### Study design

Four landscape types (urban, peri-urban, rural, and sylvatic) were selected for *Aedes* mosquito sampling in each country. The minimum distance between the sampling sites was 5 km. Landscape types are categorized following the definitions provided by Zahouli et al. (2016). Urban areas were characterized by high population densities, paved roads, and land use dominated by residential, commercial, cultural, and recreational buildings, interspersed with designated green spaces. Peri-urban sites, with lower population densities, included a mix of modern residential buildings, hospitals, schools, and ongoing urbanization in previously undeveloped areas, often with paved road infrastructure. Rural areas were characterized by low population densities, unpaved roads, a mix of traditional and modern housing, and a predominant focus on agricultural activities. Sylvatic sampling sites represented natural or minimally disturbed habitats, such as dense rainforests in Côte d’Ivoire and Ghana and wooded savannah in Burkina Faso, distinguished by dense vegetation, tree canopies, and a rich diversity of wildlife (Zahouli et al. 2016). The four landscape types, namely urban, peri-urban, rural, and sylvatic sites were represented by 1200Lgts, Goundry, Koassa and Niangolokoin in Burkina Faso, Bonoua, Samo, Koffikro and Hévéa in Côte d’Ivoire, and Godokpe, Lokoe, Kpenoe and Klefe in Ghana (Figure 1).

### Mosquito sampling and processing

*Aedes aegypti* mosquitoes were sampled between September 2021 and September 2022 (Table S1). *Aedes* mosquitoes were sampled as 4^th^ instar larvae or pupae from their aquatic habitats (e.g., tires, cans, discarded containers, water storage, tree holes) and using ovitrap in sylvatic environment to collect eggs.

Immature mosquitoes were collected using a sieve and stored separately according to sampling sites. Containers were labeled with sampling sites and dates, then transported in cool boxes to the field laboratory, where they were reared to adult under ambient environmental conditions. Freshly emerged adults were killed by freezing at −20°C and identified to species based on morphological characteristics (Becker et al. 2020). For further morphological, morphometric, and genetic analyses, at least 50 (25 female and 25 male) specimens of *Ae. aegypti* were selected per landscape type.

### Morphological analyses

#### Morphological scaling pattern analysis

Each *Ae. aegypti* specimen was visually characterized under the stereomicroscope (Leica M205 C, Leica Microsystems, Wetzlar, Germany). The scales of the first abdominal tergite of each specimen were examined and classified into five categories according to the percentage of the white scales (i.e., 0%, <20%, <40%, <60%, and <80%), as described by McBride et al. (2014). A total of 564 specimens (200 from Burkina Faso,194 from Côte d’Ivoire, and 170 from Ghana) were examined for their scaling patterns (Table S1).

#### Wing geometric morphometrics analysis

The right and left wings of each *Ae. aegypti* specimen examined for abdominal scaling were removed with fine tweezers and mounted under a cover slip with Euparal (Carl Roth, Karlsruhe, Germany). Pictures of each wing were taken at 20× magnification using a stereomicroscope (Leica M205 C, Leica Microsystems, Wetzlar, Germany). The Fiji package of ImageJ was used to digitize the coordinates of 18 landmarks on the anatomical junctions of the wing veins (Schindelin et al. 2012). The selected landmark configuration aligned with previous geometric morphometric studies in mosquito research (Lorenz et al. 2012; Sauer et al. 2020; Wilke et al. 2016). Landmarks were collected by a single observer (EA) to minimize variability and ensure measurement accuracy. A generalized Procrustes analysis was performed to superimpose the raw landmark coordinates (Adams and Otárola-Castillo 2013). The resulting Procrustes coordinates described the size-free wing shape and were used to statistically compare the wing shape variation between the countries and the landscape types. This analysis was performed using a Procrustes ANOVA with the procD.lm function (Adams and Otárola-Castillo 2013) employing 1,000 permutations. A linear discriminant analysis was conducted to visualize the wing shape variance in relation to the countries and the landscape types. The statistical analyses and visualization were performed in R (R CoreTeam 2023), including the packages geomorph (Adams and Otárola-Castillo 2013; Baken et al. 2021), MASS (Venables and Ripley 2002), and ggplot2 (Wickham 2016).

### Genetic analyses

#### Mitochondrial genome sequencing

We conducted a shotgun DNA sequencing of the *Ae. aegypti* samples. Whole single specimens were homogenized in 500 µl PBS with tungsten carbide beads (1,5 mm in diameter; Qiagen, Hilden, Germany) using TissueLyser II (Qiagen, Hilden, Germany) for 2 min at 50 Hz. After centrifugation, the DNA was extracted from the supernatant with the DNeasy Blood & Tissue Kit (Qiagen, Hilden, Germany) based on the manufacturer’s recommendation. The extracted DNA was subjected to library preparation using a QIAseq FX DNA Library Kit (Qiagen, Hilden, Germany). During the procedure, the quantity of the samples was measured with Qubit Fluorometer using Qubit DNA/RNA HS Assay Kit (Thermo Fisher Scientific, Austin, TX, USA). Final libraries were pooled together and sequenced on a NextSeq2000 machine with the usage of 200-cycle (2 × 100 bp paired-end) NextSeq2000 reagent kit (Illumina, San Diego, CA, USA).

Raw reads were quality checked and the adapter sequences were trimmed using CLC Genomics Workbench 9.0 (QIAGEN, Aarhus, Denmark). The clean reads were de novo assembled using Megahit v1.2.9 (Li et al. 2015). Reads were mapped to a reference sequence (EU352212) and mitochondrion associated contigs with Bowtie2 v2.4.5 (Langmead and Salzberg 2012). An alternative assembly method was also applied, namely the GetOrganelle v1.7.7.1 toolkit with the implementation of all available complete *Ae. aegypti* mitochondrial sequences as seed. Raw sequencing data is available in Sequence Read Archive under the Bioproject ID: PRJNA1188586. Assembled mitogenomes were uploaded to the NCBI GenBank (PQ635914-PQ636016). The sample names, Biosample IDs, and Accession numbers are linked in Supplementary Table S2.

#### Population structure analysis

Protein-coding DNA regions were extracted from the sequenced data and alignments were constructed using Geneious® 9.0.5 (Biomatters, Auckland, New Zealand). Tassel 5.2 (Bradbury et al. 2007) was used to extract SNP and create a variant call format (VCF) data file which was converted to structure format with PGDSpider 2.1 (Lischer and Excoffier 2012). Bayesian iterative algorithm implemented in STRUCTURE (Pritchard, Stephens, and Donnelly 2000) was used to analyze differences in the distribution of genetic variants among the populations. We performed 10 independent runs, with the number of cluster (K) varying from 1 to 10 at 50,000 Markov Chain Monte-Carlo (MCMC) repetitions and a burn-in period of 20,000 iterations. K was calculated using the online version of CLUMPAK (Kopelman et al. 2015).

#### Phylogenetic analysis

To determine the evolutionary relationships among the *Ae. aegypti* populations, we performed a phylogenetic analysis of the concatenated protein-coding genes (11,231 positions) on nucleotide level. The multiple sequence alignment was performed with MAFFT v7.490 (Katoh and Standley 2013). Model selection and Maximum likelihood phylogenetic tree construction was conducted with IQ-TREE 2 v2.0.7 (Minh et al. 2020). The best fitting model was selected based on the Bayesian information criterion and Akaike information criterion. Statistical support was tested using ultra-fast bootstrapping method with 1000 iterations (Minh, Nguyen, and von Haeseler 2013). The final tree was visualized and edited in TreeViewer v2.2.0 (Bianchini and Sánchez-Baracaldo 2024). Additionally, a pairwise distance matrix was generated MEGA v11.0.13 (Tamura, Stecher, and Kumar 2021). The concatenated PCGs at the nucleotide level were analyzed using Kimura 2-parameter model, including transitions and transversions and using uniform rates among sites.

We reconstructed the phylogenetic relationship between the *Ae. aegypti* samples with an ortholog based method using read2tree v0.1.5 pipeline (Dylus et al. 2024). The orthologous matrix was generated from the seven available members of *Culicidae* family, collected from OMA browser (Altenhoff et al. 2024). The sequencing depth and the sample quality was not sufficient for nuclear genome based phylogenetic analysis due to the low endogenous content.

### Morphology and genetics relationship

To assess the relationship between the *Ae. aegypti* wing shape and genetic distance, a pairwise Procrustes distance matrix was generated using the superimposed landmarks of the right wings. The obtained distance matrix based on the wing shape data was statistically compared with the pairwise genetic distance matrix using Mantel tests with Pearson’s correlation coefficient and 999 permutations (Oksanen et al. 2019). We statistically compared the two genetically distinct clusters using a Procrustes ANOVA, performed with the function procD.LM in the R package geomorph (Baken et al. 2021). This was conducted separately for the right and left wings.

## Results

### Morphological analyses

#### Morphological scaling patterns

The scaling patterns of the first abdominal tergite of Ae. aegypti populations profoundly varied between countries and landscape types (Figure 2). The highest proportion of black scales on the first tergite was found in Côte d’Ivoire (85.5%, n = 194), followed by Ghana (79.8%, n = 170) and Burkina Faso (50.4%, n = 200). Overall, the Ivorian and Ghanaian specimens had similar scaling patterns that substantially differed from those from Burkina Faso. At the sylvatic site, all specimens in Ghana were black-scaled and only one specimen had up to 20% white scales in Côte d’Ivoire. In Burkina Faso, the sylvatic populations of *Ae. aegypti* were mainly white-scaled, with over 93.3% individuals having white scales. Moreover, 42.6% urban specimens from Burkina Faso had white abdominal scales.

**Figure 2:**
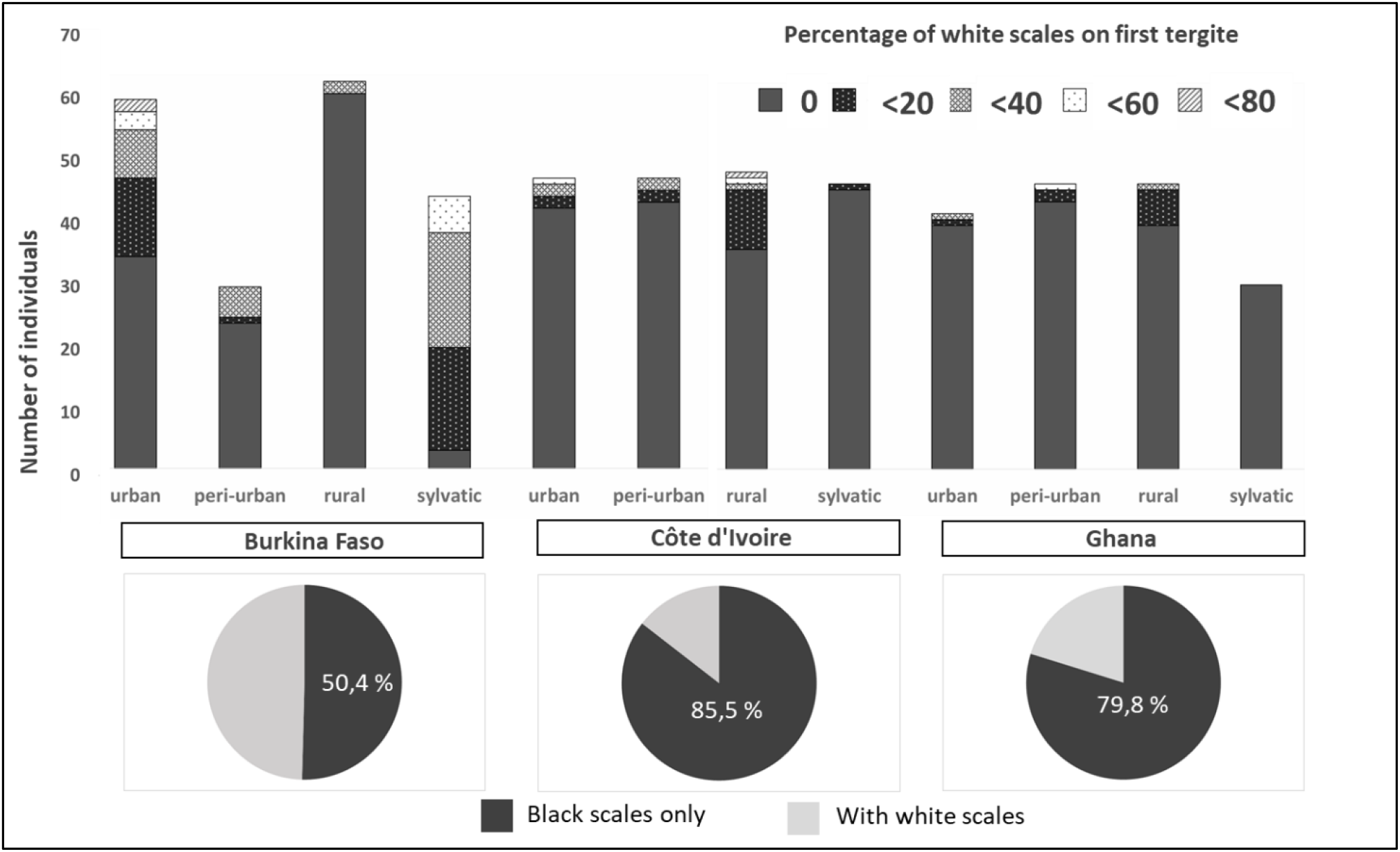
Scaling patterns (black *vs*. white) on the first abdominal tergite in *Aedes aegypti* populations per country and landscape type in Burkina Faso, Côte d’Ivoire, and Ghana.

#### Wing geometric morphometrics

Procrustes ANOVA results indicated that female wing shape was significantly influenced by the factors country and landscape type for both left (country: F = 5.02, R^2^ = 0.03, p < 0.001; landscape type: F = 3.20, R^2^ = 0.03, p < 0.001) and right (country: F = 3.76, R^2^ = 0.02, p < 0.001; landscape type: F = 2.91, R^2^ = 0.03, p < 0.001) wings. Similarly, male wing shape was significantly affected by country and landscape type for both left wings (country: F = 9.36, R^2^ = 0.07, p < 0.001; landscape type: F = 2.05, R^2^ = 0.02, p = 0.003) and right wings (country: F = 3.58, R^2^ = 0.03, p < 0.001; landscape type: F = 2.62, R^2^ = 0.03, p < 0.001). However, variances (R^2^-values) were relatively low, and linear discriminant analyses of the wing shape variation showed a distinct overlap between *Ae. aegypti* populations from different countries and different landscape types (Figure 3, A-B).

**Figure 3:**
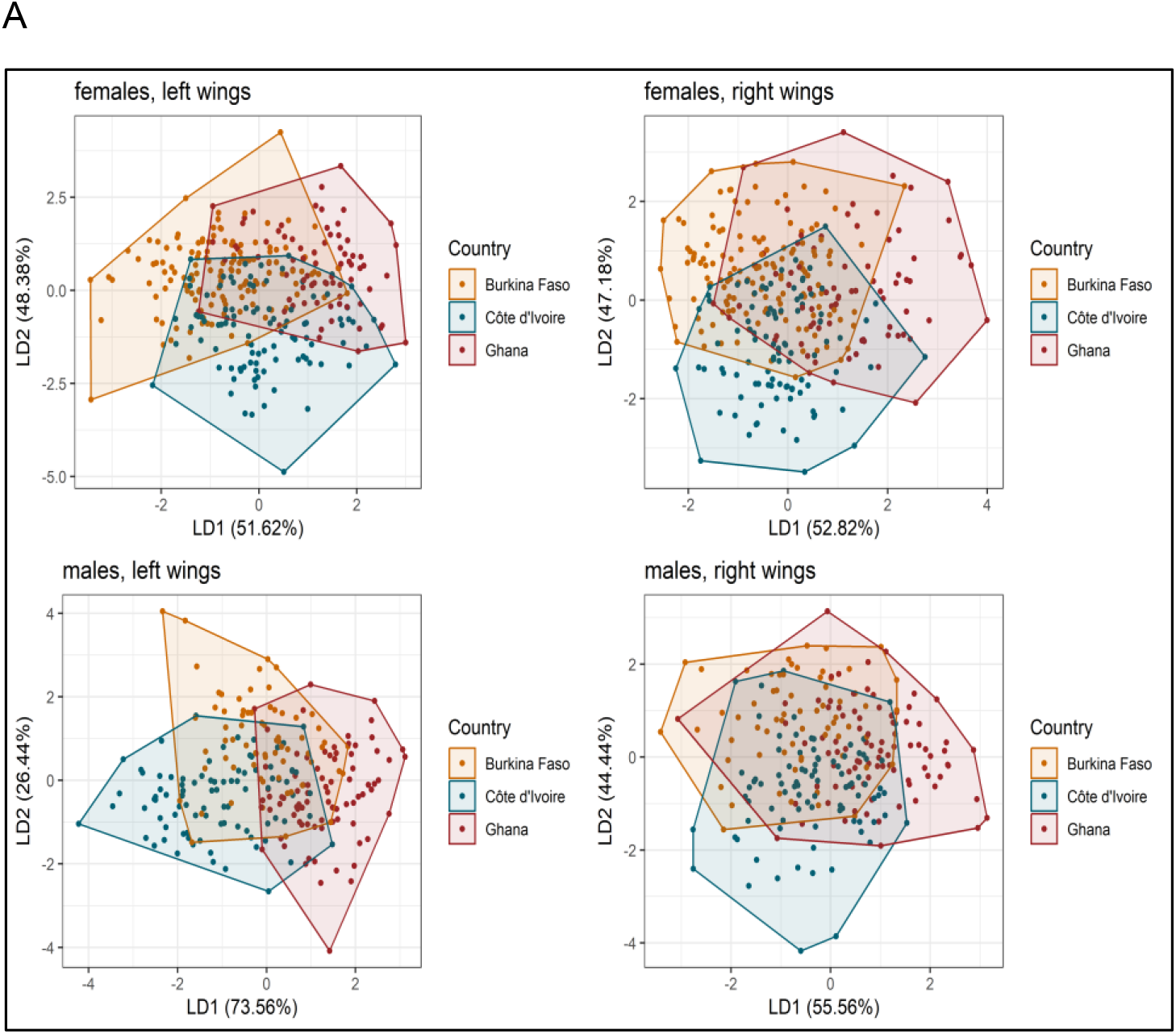
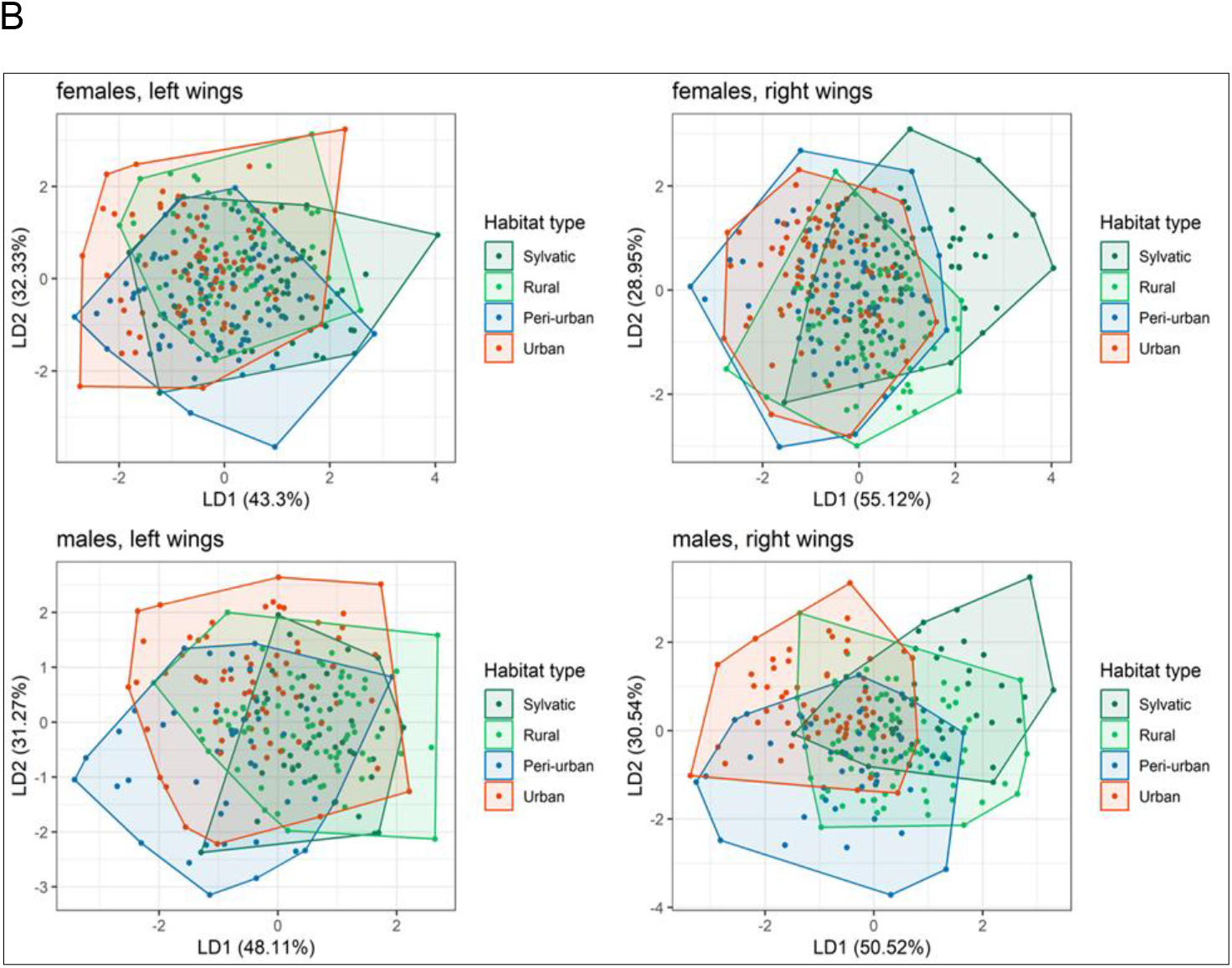
Visualization of the linear discriminant (LD) analysis showing the wing shape variation for *Aedes aegypti* from West Africa according to A) country, and B) landscape type.

### Genetic analyses

#### Population structure

The results of the STRUCTURE analysis of the samples from Burkina Faso, Côte d’Ivoire, and Ghana are shown in Figure 4. Of the 180 Ae. aegypti specimens initially selected, high-quality DNA sequences were successfully obtained for 103 (31 from Burkina Faso, 57 from Côte d’Ivoire and 15 from Ghana), which were subsequently used for genetic structure analyses. All Ae. aegypti samples clustered with two exceptions: the sylvatic specimens from Burkina Faso and a specimen from the urban site in Côte d’Ivoire. One specimen collected in the urban site and one from the rural site in Burkina Faso showed low admixture (Figure 4, K = 2). Populations from Ghana, Côte d’Ivoire, and Burkina Faso showed the same admixed pattern for K value equal to 3 (K = 3), except for those from the sylvatic site in Burkina Faso. Peri-urban populations from Burkina Faso clustered with a majority of urban populations from Côte d’Ivoire (K = 6, green). The Ivoirian and Ghanaian populations displayed a mixture of three sub clusters, including all landscape types.

**Figure 4:**
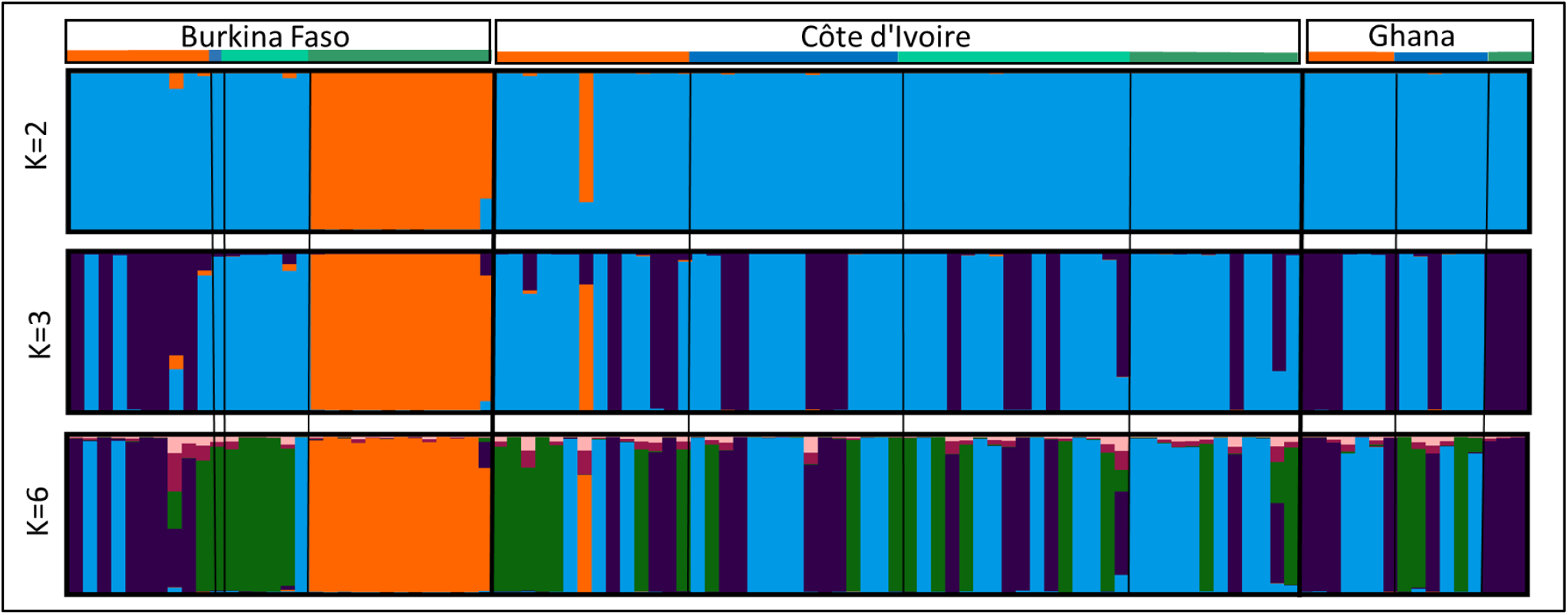
STRUCTURE bar plots for Aedes aegypti populations from Burkina Faso, Côte d’Ivoire and Ghana (West Africa) estimated using 637 SNP loci of the mitogenome with K = 2, K = 3, and K = 6. Population origins and names are reported above the plots with the colour orange referring to urban, blue to peri-urban, light green to rural and dark green to sylvatic populations. The probability of each individual assigned to a specific genetic cluster identified by STRUCTURE is depicted on the y-axis, with each group represented by distinct colours. Each bar on the graph corresponds to an individual, and those with a 100% assignment to a particular cluster are shown in a single colour. The bars display varying percentages of different colours for individuals with mixed ancestry.

#### Phylogenetics

The phylogeny of protein-coding genes revealed the presence of two major mitochondrial lineages A and B (Figure 5). The majority of samples belong to clade B showed no clustering by country or landscape type. One sample collected in urban sites from Burkina Faso (252) is forming a single branch in clade B. Specimens from the sylvatic site in Burkina Faso clustered in clade A together with an urban sample in Côte d’Ivoire (432) and a sylvatic sample in Burkina Faso (351).

**Figure 5:**
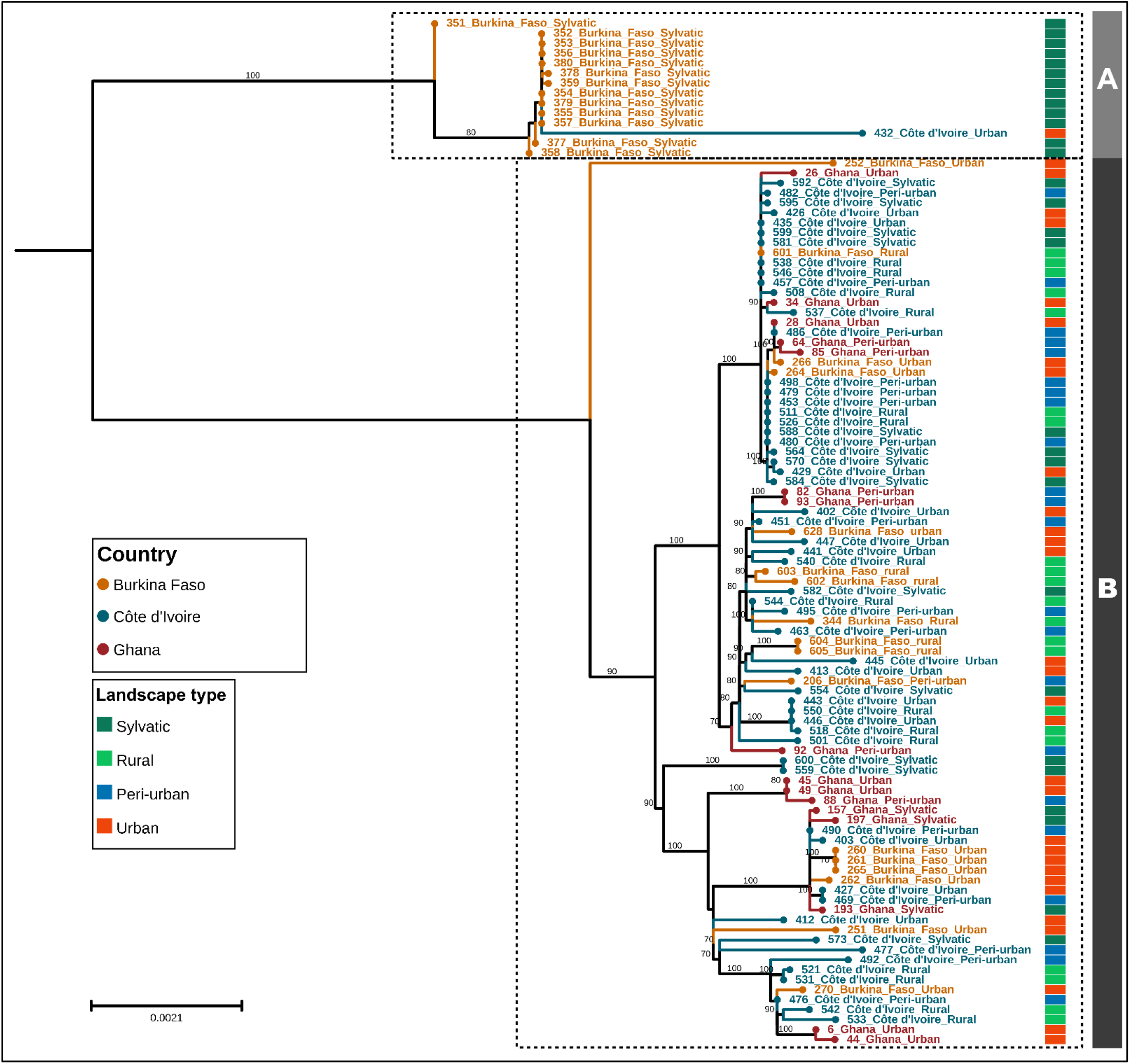
Mid-point rooted, Maximum likelihood (ML) phylogenetic tree constructed using13 PCGs. HKY+F+R2 substitution model was used based on the model selection with 1000 UltraFast bootstrap. Geographical and landscape associated annotations were added to the tree.

### Morphology and genetics relationship

The Mantel tests revealed no significant correlations between the wing shapes and genetic distances for either females (r = −0.063, p = 0.785) or males (r = 0.050, p = 0.22). The wing shape of the females did not differ significantly between the two genetically distinct clusters (Figure 5, A-B) for either the left (df = 67, F = 0.907, R^2^ = 0.014, p = 0.494) or the right (df = 64, F = 1.205, R^2^ = 0.0188, p = 0.263) wings (Figure S1). No results were obtained for males due to an insufficient number of successfully sequenced specimens for statistical analysis. The generated heatmap visualizing the relationship between wing shape and genetic distance confirmed the absence of a significant correlation between them (Figure S2).

## Discussion

### Morphology

The *Ae. aegypti* populations studied revealed white and black scales on the first abdominal tergite with considerable variations observed across different countries and landscape types. The studied populations from Côte d’Ivoire and Ghana exhibited comparable scaling patterns. In marked contrast, the sylvatic populations in Burkina Faso displayed a distinct scaling pattern. Rose et al. (2020) demonstrated that abdominal scaling was strongly correlated with preference for humans across Africa, especially in Burkina Faso. These divergences found between countries and landscape types provide critical insights into regional variation in the morphology of *Ae. aegypti* populations (Rose et al. 2023). These morphological variations may have resulted from different environmental pressures, such as abiotic factors, resource availability that might influence the bio-ecological characteristics of the local populations (Ouédraogo et al. 2022). Geographical variation, localized evolutionary processes, genetic drift, and climatic conditions may impose selective pressures on shaping morphological features and contribute to phenotypic plasticity (Bennett, McMillan, and Loaiza 2021; Britos Molinas, Gayozo Melgarejo, and Rojas de Arias 2022). In Senegal, studies have demonstrated significant variations in abdominal pale scale patterns, with both forms often coexisting within the same populations (Diouf et al. 2020; Paupy et al. 2010). Moreover, temporal variations in scaling patterns within the Aaa form have been identified, with mosquitoes segregating into two primary morphological groups influenced by seasonal fluctuations and breeding site characteristics (Paupy et al. 2010). The traditional classification of *Ae. aegypti* into the two subspecies (Aaa and Aaf) was questioned. Notable morphological differences among populations were found, casting doubt on the validity of this subspecies distinction, mainly in West Africa (Diouf et al. 2020). However, it has been shown that Aaa and Aaf are genetically distinct (Gloria-Soria et al. 2016). Due to these disparities, subspecies designation has become complex.

The wing shape analysis showed a clustering according to country and landscape type, but the variance was low. Previous studies have demonstrated the utility of wing morphometric analysis for distinguishing *Ae. aegypti* populations across different environmental and geographical contexts. For example, Wilk-da-Silva et al. (2018) identified distinct wing morphometric patterns among *Ae. aegypti* populations from areas with varying levels of urbanization, achieving high reclassification accuracy of individuals to their respective populations. A recent study in Benin (West Africa) showed morphometric divergences in *Ae. aegypti* populations across various locations, including urban, semi-urban, and sylvatic sites (Hounkanrin et al. 2023). Similarly, other studies reported significant differences in wing shape between *Ae. aegypti* populations from different geographical regions (Carvajal et al. 2016; Chaiphongpachara, Juijayen, and Khlaeo Chansukh 2019; Leyton Ramos et al. 2020). Studies reported that *Ae. aegypti* wing size was positively associated with breeding container types, and water physicochemical parameters such as electrical conductivity, and negatively with pH and temperature and with the level of exposure of the container to sunlight (Ouédraogo et al. 2022). While these abiotic factors influence wing size in mosquitoes, wing shape provides insights into heritable intraspecific and geographic differences (Dujardin 2008; Henry et al. 2010; Morales Vargas et al. 2013). This justifies the use of wing shape and the silence on wing size in our current study. Differences in seasonal conditions at sampling sites and relatively small sample sizes could affect the robustness of our findings and other studies. Future studies should incorporate larger sample sizes and standardized sampling across multiple seasons to better understand the interaction between environmental variables, landscape types, and mosquito morphology.

Wing shape analysis can be a valuable tool for assessing mosquito population structure, but its success depends on the context. In some studies, wing shape analysis successfully differentiated populations, while in others, the results were less conclusive (Francuski et al. 2016; Hounkanrin et al. 2023). In our study, the slightly significant results should be interpreted with caution. The lack of strong differentiation between populations, even across considerable geographical distances, suggests either substantial gene flow between populations or limited sensitivity of the method to detect subtle differences in *Ae. aegypti* populations from our study regions. The results concerning the sylvatic populations in Burkina Faso, for instance, did not show a high level of differentiation, further supporting this observation.

### Genetics

Among the 103 individuals analyzed, most specimens showed substantial genetic mixing, a high degree of gene flow among populations across the study regions. However, two notable outliers were observed: the sylvatic samples from Burkina Faso and one urban sample from Côte d’Ivoire (432), which did not cluster with the overall populations of the focus countries, suggesting distinct genetic lineages.

Admixture analysis at K = 2 indicated that most West African populations share a common genetic background except the sylvatic samples from Burkina Faso and one urban sample from Côte d’Ivoire. Populations from Ghana, Côte d’Ivoire are showing the same admixed pattern at K = 3. A study by Abuelmaali et al. (2024) identified a notable correlation between genetic variations and geographical distance. This correlation implies that gene flow and migration among populations are significantly influenced by their proximity in geographical terms (Abuelmaali et al. 2024). However, in our sampled populations, no clear correlation between genetic variation and geographical distance was observed, suggesting additional factors may be influencing genetic structure in these regions. Another study indicated that *Ae. aegypti* populations specifically from Goudiry in Senegal and Sedihou in Angola, respectively, showed signs of admixture (Kotsakiozi et al. 2018). This suggests that these populations may have experienced gene flow from other populations, possibly due to human-assisted migrations or ecological changes (Kotsakiozi et al. 2018).

The phylogenetic analysis of mitochondrial protein-coding genes identified two primary mitochondrial lineages, designated as clades A and B. This finding suggests significant genetic diversity within the *Ae. aegypti* populations across the sampled regions. Clade B encompassed the majority of specimens, showing no apparent clustering by country or landscape type, indicating a lack of strong geographic or ecological variation within this lineage. This pattern is consistent with widespread gene flow and admixture among *Ae. aegypti* populations, likely facilitated by their mobility and adaptation to diverse environments. An intriguing observation is the unique placement of the Burkina Faso urban sample (252) as a distinct branch within clade B. The outlier might represent a lineage with a distinctive evolutionary history or a recent mutation event. A report states that mutations occurring in a lineage can lead to distinctive traits that set it apart from others (Baum 2008). In contrast, clade A showed notable clustering of sylvatic specimens from Burkina Faso alongside two other samples, Côte d’Ivoire urban (432) and Burkina Faso sylvatic (351). This clustering suggests that these populations share a more recent common ancestor and may represent a lineage that is less influenced by a likely gene flow. The inclusion of an urban sample from Côte d’Ivoire within clade A could indicate historical migration events or incomplete lineage sorting rather than strict habitat-specific structuring.

### Morphology and genetics relationship

The results indicate that *Ae. aegypti* populations did not show a significant clustering of their wing shape corresponding to the identified genetic clusters, suggesting that genetic distance was not reflected in shape distance. The findings of this study aligned with broader research, implying that the wing shape in mosquitoes is influenced by a combination of genetic and environmental factors, with its utility for population structure and evolutionary studies varying across species (Evans et al. 2018; Henry et al. 2010; Jirakanjanakit et al. 2007; Morales Vargas et al. 2013; Phanitchat et al. 2019; Roux et al. 2015). Previous research on *Ae. aegypti* has indicated that wing shape exhibits a limited capacity for distinguishing populations, potentially due to stabilizing selective pressures (Vidal and Suesdek 2012). While wing shape remains a valuable morphological trait for specific evolutionary and ecological studies, its applicability for delineating population structure or genetic clusters may be constrained in certain mosquito species, including *Ae. aegypti*. Further research integrating environmental variables and additional genetic markers may help disentangle these relationships.

## Conclusion

This study provides insights into the morphological and genetic diversity of *Ae. aegypti* populations in West Africa, highlighting the complex interplay between environmental, genetic, and ecological factors. The observed variation in abdominal tergite scale patterns and wing morphometrics across regions underscores the influence of local environmental pressures and historical evolutionary processes. While scale patterns align broadly with previous findings, such as the predominance of black scales in sylvatic populations and mixed scaling patterns in urban populations, the distinct patterns observed in Burkina Faso suggest unique regional evolutionary or ecological pressures. The lack of strong differentiation in wing shape between genetic clusters suggests that wing morphology may not be a reliable marker of genetic divergence in *Ae. aegypti*. Genetic analyses revealed substantial gene flow and admixture across populations, with notable exceptions in sylvatic populations from Burkina Faso, which exhibited distinct clustering patterns. These genetic outliers suggest localized evolutionary dynamics or historical migration events, potentially linked to human-mediated dispersal.

Overall, this study underscores the importance of integrating genetic, morphological, and ecological data to unravel the evolutionary dynamics of *Ae. aegypti*. The unexpected findings in scaling patterns and genetic clustering in Burkina Faso highlight the need for further investigation to better understand how environmental and historical factors shape the phenotypic and genetic diversity of this species. These insights are crucial for refining *Aedes* vector control strategies and predicting and preventing the potential spread of mosquito-borne arboviral diseases in West Africa.

## Acknowledgment

We are grateful to the technicians, Alexander Bialonski and Konstantin Kliemke, for their technical assistance. This study was supported by the German Research Foundation (JO 1276/5-1) and the Federal Ministry of Education and Research of Germany (BMBF) under project NEED (01Kl2022).

## Data Accessibility and Benefit-Sharing Section

### Data Accessibility Statement

#### Genetic data

The nucleotide sequences of the mtDNA are available in the NCBI Genbank (Accession # PQ635914 - PQ636016).

#### Sample metadata

Details of the samples are archived under BioProject ID PRJNA1188586.

Comprehensive mosquito wing images can be found in the Bioimage Archive published under a CC-BY 4.0 license (S-BIAD1478, https://doi.org/10.6019/S-BIAD1478).

### Benefit-Sharing Statement

The benefits of this research stem from sharing our data and results through public databases. Collaboration, cooperation, and contribution in education and training were established, and scientists from other countries provided the samples. The authors of this scientific paper include all the collaborators who contributed in diverse ways. In the advent of strengthening capacity building, knowledge and technology were transferred to the provider of the genetic resources (samples) under fair, concessional, and preferential terms.

## Authors’ contribution

Conceptualization: EA, JZBZ, AB, HJ, RL, DC, JSC; data collection: EA, YNB, CNA, AS; data analysis: EA, FGS, GET, BH, DC, HJ; first drafting: EA, JZBZ, AB, HJ, AT; writing and editing: EA, JZBZ, AB, AT, JSC, RL, FGS, GET, HJ.

## Supplementary materials

**Table S1**. Population information of sampled *Aedes aegypti* mosquitoes.

**Table S2:** Summary of uploaded sequences at NCBI for BioProject PRJNA1188586.

**Figure S1:** PCA plots on the wing shape using the females belonging to the two identified genetic clusters.

**Figure S2**: Heatmap showing the differences between wing shape and genetic distance.

