## supplemental files for "Morphological and genetic heterogeneity in *Aedes aegypti* (Diptera: Culicidae) populations across diverse landscapes in West Africa"

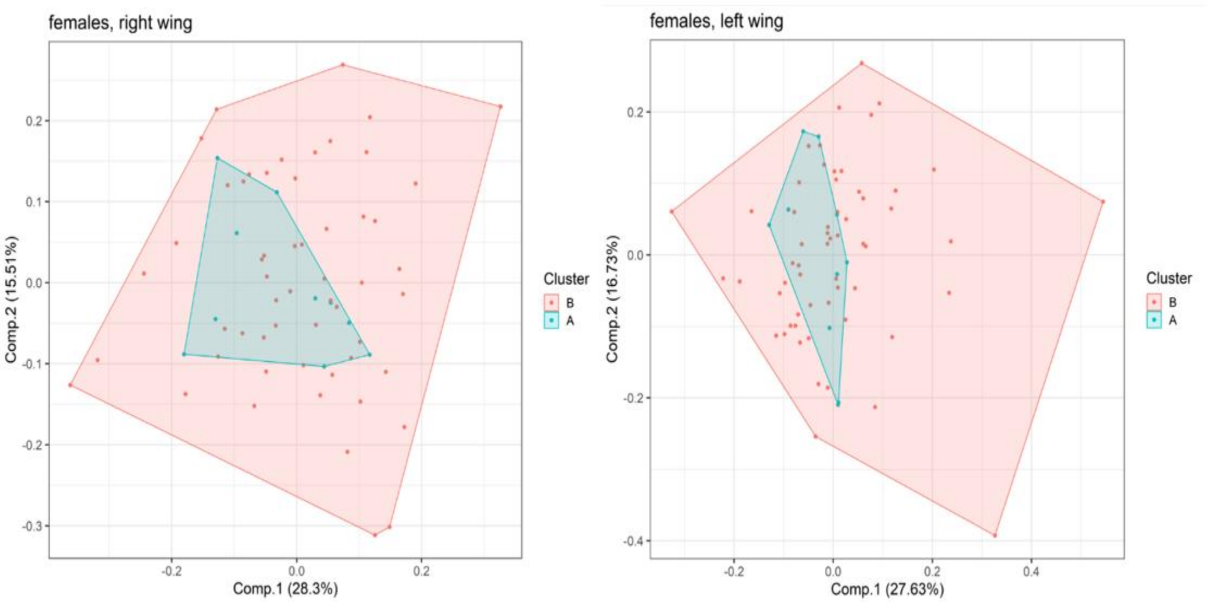


Figure S1: PCA plots on the wing shape using the females belonging to the two identified genetic clusters (A and B)

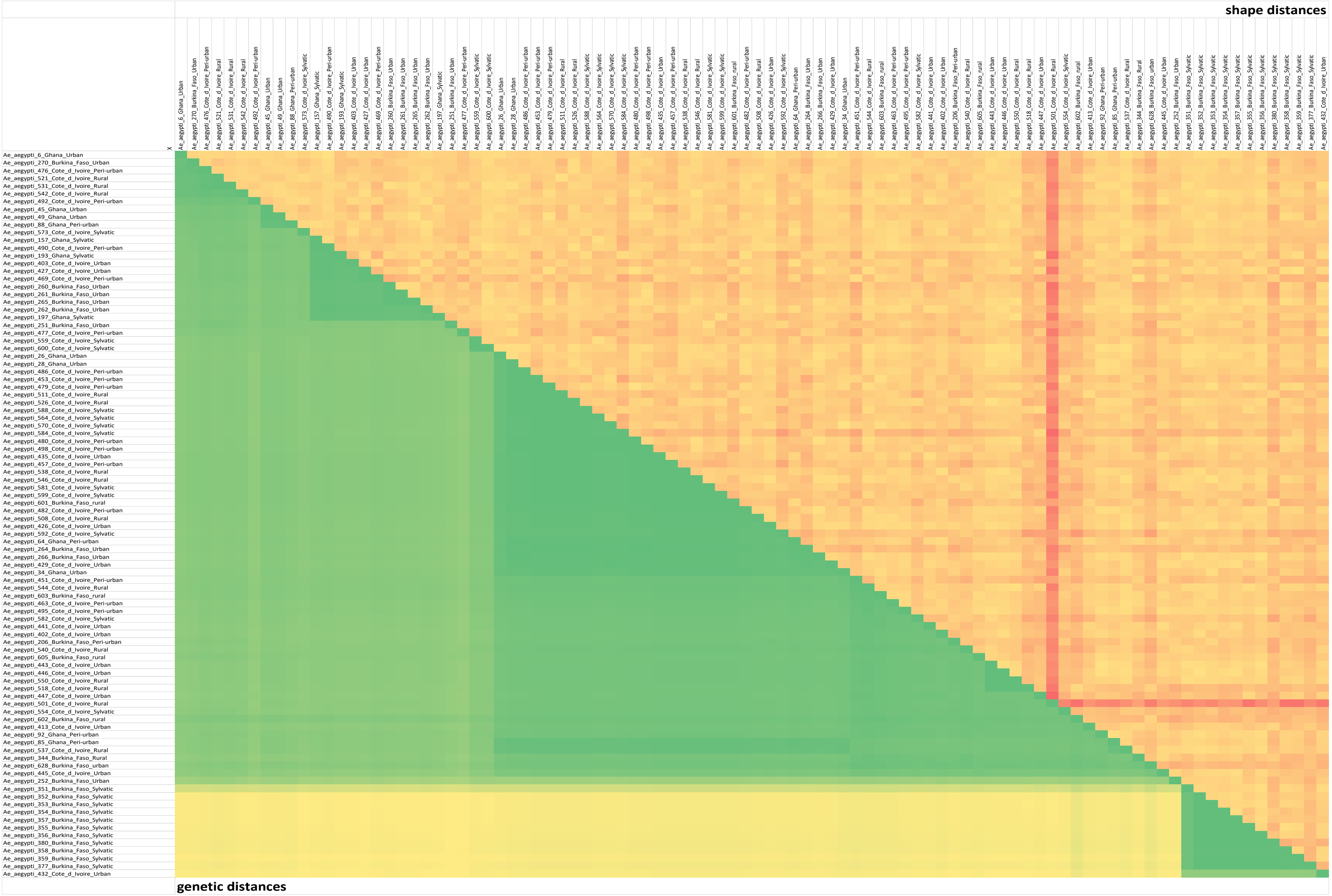


Figure S2: Heatmap showing the difference between wing shape and genetic distance.

Table S1: Population information of sampled *Aedes aegypti* mosquitoes

| Country | Sampling Period | Locality | Landscape type | Latitude | Longitude | Samples | Used for scaling pattern | Almost complete mitochondrial genome | Used for morphometric analysis (left/right wing) |
|---|---|---|---|---|---|---|---|---|---|
| **Côte d'Ivoire** | **19.07.2022 - 25.07.2022** | **Bonoua** | **urban** | **5.26299** | **-3.60133** | **50** | **48** | **14** | **48/47** |
| | | **Samo** | **peri-urban** | **5.283013** | **-3.515762** | **50** | **48** | **15** | **48/48** |
| | | **Koffikro** | **rural** | **5.404234** | **-3.378681** | **50** | **50** | **15** | **50/49** |
| | | **Hévéa** | **sylvatic** | **5.415712** | **-3.49393** | **50** | **48** | **13** | **42/44** |
| **Burkina Faso** | **18.09.2021 - 23.09.2022** | **Ouagadougou** | **urban** | **12.3741667** | **-1.50028** | **75** | **61** | **10** | **67/66** |
| | | **Goundry** | **peri-urban** | **12.517910** | **-1.341019** | **50** | **30** | **1** | **41/40** |
| | | **Koassa** | **rural** | **12.005175** | **-1.330274** | **75** | **64** | **6** | **61/63** |
| | | **Niangoloko** | **sylvatic** | **10.26562** | **-4.91394** | **50** | **45** | **13** | **33/35** |
| **Ghana** | **05.10.2021 – 30.10.2021** | **Godokpoe** | **urban** | **6.605.838** | **0.496095** | **50** | **43** | **7** | **47/46** |

| | | | | | | | | | |
|---|---|---|---|---|---|---|---|---|---|
| | | **Lokoe-Site** | **peri-urban** | **6.586.111** | **0.437192** | **50** | **48** | **6** | **45/46** |
| | | **Kpenoe** | **rural** | **6.631.574** | **0.515839** | **50** | **48** | **0** | **47/47** |
| | | **Klefe** | **sylvatic** | **6.619.748** | **0.443849** | **50** | **31** | **3** | **28/30** |
| **Total** | | | | | | **650** | **564** | **103** | **557/561** |

Table S2_Summary of uploaded sequences at NCBI for BioProject PRJNA1188586

| Sample_ID | BioProject_ID | Accession_number | Biosample_ID | Sequencing_depth | Mean_coverage |
|---|---|---|---|---|---|
| Ae.aegypti_006 | PRJNA1188586 | PQ635914 | SAMN44843032 | 7315660 | 29.1 |
| Ae.aegypti_026 | PRJNA1188586 | PQ635915 | SAMN44843033 | 8811764 | 14.1 |
| Ae.aegypti_028 | PRJNA1188586 | PQ635916 | SAMN44843034 | 6332848 | 28.4 |
| Ae.aegypti_034 | PRJNA1188586 | PQ635917 | SAMN44843035 | 8018772 | 11 |
| Ae.aegypti_044 | PRJNA1188586 | PQ635918 | SAMN44843036 | 6304410 | 25.8 |
| Ae.aegypti_045 | PRJNA1188586 | PQ635919 | SAMN44843037 | 4252818 | 15.9 |
| Ae.aegypti_049 | PRJNA1188586 | PQ635920 | SAMN44843038 | 6132690 | 43.8 |
| Ae.aegypti_064 | PRJNA1188586 | PQ635921 | SAMN44843039 | 4622292 | 11 |
| Ae.aegypti_082 | PRJNA1188586 | PQ635922 | SAMN44843040 | 7199464 | 44.9 |
| Ae.aegypti_085 | PRJNA1188586 | PQ635923 | SAMN44843041 | 6833420 | 10.7 |
| Ae.aegypti_088 | PRJNA1188586 | PQ635924 | SAMN44843042 | 5628766 | 10 |
| Ae.aegypti_092 | PRJNA1188586 | PQ635925 | SAMN44843043 | 6366286 | 8.8 |
| Ae.aegypti_093 | PRJNA1188586 | PQ635926 | SAMN44843044 | 3540286 | 166.9 |
| Ae.aegypti_157 | PRJNA1188586 | PQ635927 | SAMN44843045 | 3205994 | 53.6 |
| Ae.aegypti_193 | PRJNA1188586 | PQ635928 | SAMN44843046 | 4085414 | 14.8 |
| Ae.aegypti_197 | PRJNA1188586 | PQ635929 | SAMN44843047 | 6300414 | 16.9 |
| Ae.aegypti_206 | PRJNA1188586 | PQ635930 | SAMN44843048 | 4252258 | 94.4 |
| Ae.aegypti_251 | PRJNA1188586 | PQ635931 | SAMN44843049 | 4499710 | 39.5 |

| | | | | | |
|---|---|---|---|---|---|
| Ae.aegypti_252 | PRJNA1188586 | PQ635932 | SAMN44843050 | 5714540 | 27.6 |
| Ae.aegypti_260 | PRJNA1188586 | PQ635933 | SAMN44843051 | 6842718 | 16.4 |
| Ae.aegypti_261 | PRJNA1188586 | PQ635934 | SAMN44843052 | 4474988 | 48.9 |
| Ae.aegypti_262 | PRJNA1188586 | PQ635935 | SAMN44843053 | 4220954 | 28.1 |
| Ae.aegypti_264 | PRJNA1188586 | PQ635936 | SAMN44843054 | 4509032 | 14.5 |
| Ae.aegypti_265 | PRJNA1188586 | PQ635937 | SAMN44843055 | 4688254 | 27.7 |
| Ae.aegypti_266 | PRJNA1188586 | PQ635938 | SAMN44843056 | 4508534 | 49 |
| Ae.aegypti_270 | PRJNA1188586 | PQ635939 | SAMN44843057 | 5723204 | 24 |
| Ae.aegypti_344 | PRJNA1188586 | PQ635940 | SAMN44843058 | 5912558 | 10.1 |
| Ae.aegypti_351 | PRJNA1188586 | PQ635941 | SAMN44843116 | 6713950 | 8.6 |
| Ae.aegypti_352 | PRJNA1188586 | PQ635942 | SAMN44843117 | 3724920 | 18.2 |
| Ae.aegypti_353 | PRJNA1188586 | PQ635943 | SAMN44843118 | 4224672 | 13.5 |
| Ae.aegypti_354 | PRJNA1188586 | PQ635944 | SAMN44843119 | 3975702 | 26.3 |
| Ae.aegypti_355 | PRJNA1188586 | PQ635945 | SAMN44843120 | 5177824 | 19.1 |
| Ae.aegypti_356 | PRJNA1188586 | PQ635946 | SAMN44843121 | 5310424 | 18.9 |
| Ae.aegypti_357 | PRJNA1188586 | PQ635947 | SAMN44843122 | 5839444 | 27.6 |
| Ae.aegypti_358 | PRJNA1188586 | PQ635948 | SAMN44843123 | 5207254 | 12.5 |
| Ae.aegypti_359 | PRJNA1188586 | PQ635949 | SAMN44843124 | 3066320 | 9.2 |
| Ae.aegypti_377 | PRJNA1188586 | PQ635950 | SAMN44843125 | 6348552 | 8.5 |
| Ae.aegypti_378 | PRJNA1188586 | PQ635951 | SAMN44843126 | 5215952 | 11.7 |

| | | | | | |
|---|---|---|---|---|---|
| Ae.aegypti_379 | PRJNA1188586 | PQ635952 | SAMN44843127 | 5971360 | 17.2 |
| Ae.aegypti_380 | PRJNA1188586 | PQ635953 | SAMN44843128 | 7207998 | 13.9 |
| Ae.aegypti_402 | PRJNA1188586 | PQ635954 | SAMN44843059 | 7822594 | 26.7 |
| Ae.aegypti_403 | PRJNA1188586 | PQ635955 | SAMN44843060 | 7071014 | 49.9 |
| Ae.aegypti_412 | PRJNA1188586 | PQ635956 | SAMN44843061 | 5329844 | 53.2 |
| Ae.aegypti_413 | PRJNA1188586 | PQ635957 | SAMN44843062 | 5550174 | 16.1 |
| Ae.aegypti_426 | PRJNA1188586 | PQ635958 | SAMN44843063 | 6745534 | 20.9 |
| Ae.aegypti_427 | PRJNA1188586 | PQ635959 | SAMN44843064 | 4812266 | 34 |
| Ae.aegypti_429 | PRJNA1188586 | PQ635960 | SAMN44843065 | 3145194 | 10.8 |
| Ae.aegypti_432 | PRJNA1188586 | PQ635961 | SAMN44843066 | 3216832 | 22.1 |
| Ae.aegypti_435 | PRJNA1188586 | PQ635962 | SAMN44843067 | 4840600 | 25.9 |
| Ae.aegypti_441 | PRJNA1188586 | PQ635963 | SAMN44843068 | 4198840 | 30.8 |
| Ae.aegypti_443 | PRJNA1188586 | PQ635964 | SAMN44843069 | 6506300 | 76.4 |
| Ae.aegypti_445 | PRJNA1188586 | PQ635965 | SAMN44843070 | 4382470 | 7.1 |
| Ae.aegypti_446 | PRJNA1188586 | PQ635966 | SAMN44843071 | 3311332 | 58.6 |
| Ae.aegypti_447 | PRJNA1188586 | PQ635967 | SAMN44843072 | 3754422 | 24.8 |
| Ae.aegypti_451 | PRJNA1188586 | PQ635968 | SAMN44843073 | 3335106 | 26.6 |
| Ae.aegypti_453 | PRJNA1188586 | PQ635969 | SAMN44843074 | 2904432 | 29.5 |
| Ae.aegypti_457 | PRJNA1188586 | PQ635970 | SAMN44843075 | 4035532 | 51.2 |
| Ae.aegypti_463 | PRJNA1188586 | PQ635971 | SAMN44843076 | 6448652 | 33.3 |

| | | | | | |
|---|---|---|---|---|---|
| Ae.aegypti_469 | PRJNA1188586 | PQ635972 | SAMN44843077 | 3672518 | 12.6 |
| Ae.aegypti_476 | PRJNA1188586 | PQ635973 | SAMN44843078 | 3402980 | 57.2 |
| Ae.aegypti_477 | PRJNA1188586 | PQ635974 | SAMN44843079 | 1776608 | 49 |
| Ae.aegypti_479 | PRJNA1188586 | PQ635975 | SAMN44843080 | 3266382 | 35 |
| Ae.aegypti_480 | PRJNA1188586 | PQ635976 | SAMN44843081 | 2773490 | 40.3 |
| Ae.aegypti_482 | PRJNA1188586 | PQ635977 | SAMN44843082 | 2902088 | 45.4 |
| Ae.aegypti_486 | PRJNA1188586 | PQ635978 | SAMN44843083 | 5668582 | 37.9 |
| Ae.aegypti_490 | PRJNA1188586 | PQ635979 | SAMN44843084 | 5596862 | 51.3 |
| Ae.aegypti_492 | PRJNA1188586 | PQ635980 | SAMN44843085 | 5467404 | 95.2 |
| Ae.aegypti_495 | PRJNA1188586 | PQ635981 | SAMN44843086 | 3367744 | 21.8 |
| Ae.aegypti_498 | PRJNA1188586 | PQ635982 | SAMN44843087 | 5944886 | 81.6 |
| Ae.aegypti_501 | PRJNA1188586 | PQ635983 | SAMN44843088 | 4832200 | 16.6 |
| Ae.aegypti_508 | PRJNA1188586 | PQ635984 | SAMN44843089 | 6614794 | 102.7 |
| Ae.aegypti_511 | PRJNA1188586 | PQ635985 | SAMN44843090 | 5339314 | 23.6 |
| Ae.aegypti_518 | PRJNA1188586 | PQ635986 | SAMN44843091 | 6890722 | 38.2 |
| Ae.aegypti_521 | PRJNA1188586 | PQ635987 | SAMN44843092 | 5713810 | 31.9 |
| Ae.aegypti_526 | PRJNA1188586 | PQ635988 | SAMN44843093 | 7486644 | 58.4 |
| Ae.aegypti_531 | PRJNA1188586 | PQ635989 | SAMN44843094 | 5494982 | 58.2 |
| Ae.aegypti_533 | PRJNA1188586 | PQ635990 | SAMN44843095 | 6735848 | 47.7 |
| Ae.aegypti_537 | PRJNA1188586 | PQ635991 | SAMN44843096 | 6538712 | 20 |

| | | | | | |
|---|---|---|---|---|---|
| Ae.aegypti_538 | PRJNA1188586 | PQ635992 | SAMN44843097 | 6253276 | 45 |
| Ae.aegypti_540 | PRJNA1188586 | PQ635993 | SAMN44843098 | 5383724 | 29.4 |
| Ae.aegypti_542 | PRJNA1188586 | PQ635994 | SAMN44843099 | 5487140 | 35.1 |
| Ae.aegypti_544 | PRJNA1188586 | PQ635995 | SAMN44843100 | 6589052 | 59.5 |
| Ae.aegypti_546 | PRJNA1188586 | PQ635996 | SAMN44843101 | 6190852 | 43.2 |
| Ae.aegypti_550 | PRJNA1188586 | PQ635997 | SAMN44843102 | 8692338 | 75.1 |
| Ae.aegypti_554 | PRJNA1188586 | PQ635998 | SAMN44843103 | 7005782 | 43.9 |
| Ae.aegypti_559 | PRJNA1188586 | PQ635999 | SAMN44843104 | 12312700 | 104.9 |
| Ae.aegypti_564 | PRJNA1188586 | PQ636000 | SAMN44843105 | 4601716 | 85.1 |
| Ae.aegypti_570 | PRJNA1188586 | PQ636001 | SAMN44843106 | 5775666 | 36.5 |
| Ae.aegypti_573 | PRJNA1188586 | PQ636002 | SAMN44843107 | 5530106 | 29.1 |
| Ae.aegypti_581 | PRJNA1188586 | PQ636003 | SAMN44843108 | 7137744 | 76.8 |
| Ae.aegypti_582 | PRJNA1188586 | PQ636004 | SAMN44843109 | 8978866 | 64.2 |
| Ae.aegypti_584 | PRJNA1188586 | PQ636005 | SAMN44843110 | 8573222 | 32.8 |
| Ae.aegypti_588 | PRJNA1188586 | PQ636006 | SAMN44843111 | 7967914 | 59.8 |
| Ae.aegypti_592 | PRJNA1188586 | PQ636007 | SAMN44843112 | 4737372 | 21.5 |
| Ae.aegypti_595 | PRJNA1188586 | PQ636008 | SAMN44843113 | 5121302 | 45.2 |
| Ae.aegypti_599 | PRJNA1188586 | PQ636009 | SAMN44843114 | 5187630 | 36.5 |
| Ae.aegypti_600 | PRJNA1188586 | PQ636010 | SAMN44843115 | 5840786 | 80.5 |
| Ae.aegypti_601 | PRJNA1188586 | PQ636011 | SAMN44843129 | 5712272 | 22.9 |

| | | | | | |
|---|---|---|---|---|---|
| Ae.aegypti_602 | PRJNA1188586 | PQ636012 | SAMN44843130 | 4445194 | 8.8 |
| Ae.aegypti_603 | PRJNA1188586 | PQ636013 | SAMN44843131 | 6040972 | 117.7 |
| Ae.aegypti_604 | PRJNA1188586 | PQ636014 | SAMN44843132 | 5868288 | 45.4 |
| Ae.aegypti_605 | PRJNA1188586 | PQ636015 | SAMN44843133 | 5778108 | 40.6 |
| Ae.aegypti_628 | PRJNA1188586 | PQ636016 | SAMN44843134 | 5089552 | 6.9 |
